# Systematic identification and characterization of virus lncRNAs suggests extensive structural mimicry of host lncRNAs

**DOI:** 10.1101/2024.11.08.622600

**Authors:** Ping Fu, Zena Cai, Ruina You, Lei Deng, Zhaoyong Li, Zhichao Miao, Yousong Peng

## Abstract

Virus long non-coding RNAs (vlncRNAs) play crucial roles in viral infections, yet their identification and characterization remain limited. This study identified 5,053 novel vlncRNAs across 25 viral species using third-generation sequencing, with two from Influenza A virus and Vesicular stomatitis virus validated by RT-qPCR. Most vlncRNAs originated from dsDNA viruses. Only 1% having annotated RNA families, suggesting many novel RNA structures. Although only seven vlncRNAs shared sequence similarities with human lncRNAs (hlncRNAs), 772 vlncRNAs from 15 human viruses structurally mimicked hlncRNAs. These vlncRNA and hlncRNAs bound to similar miRNAs, potentially acting as miRNA sponges to promote essential life process. Splicing analysis showed vlncRNAs had a prevalence of alternative first exon. Finally, we developed vlncRNAbase (http://computationalbiology.cn/vlncRNAbase/#/) to store and organize the newly identified and known vlncRNAs. Overall, the study provides a valuable resource for further investigation into vlncRNAs and deepens our understanding of the diversity, structure and function of the molecule.

## Introduction

The long non-coding RNA (lncRNA) is a prominent class of non-coding transcripts that have longer than 200 nt and lack the protein-coding ability^[1]^. The majority of lncRNAs are transcribed by the RNA polymerase II, with typical 5’ capping and 3’ polyadenylated tailing^[2]^. The lncRNAs have been widely found in humans, animals, plants, fungi and prokaryotes^[3]^. They have diverse functions, including modulating gene expression^[4]^, involving in chromatin structure^[5]^ and contributing to cell differentiation^[6]^. The lncRNAs play an important role in multiple human diseases, including cancer^[7–9]^. For example, the lncRNA PVT1 was reported to promote ovarian cancer progression by silencing miR-214, a miRNA commonly involved in carcinogenesis^[7]^. Additionally, the lncRNA PNUTS was involved in breast cancer metastasis through its impact on the epithelial-mesenchymal transition process^[8]^. Thus, they have been considered crucial components of the complex regulatory network involved in disease development^[9]^.

The rapid development of high-throughput sequencing technology has greatly facilitated the identification and characterization of lncRNAs^[10,11]^. Numerous lncRNAs have been identified based on the next-generation-sequencing (NGS) technology, and several lncRNA databases have been built^[12–14]^. For example, the lncRNAfunc database contains 15,900 human lncRNAs that are involved in 33 cancer types^[12]^ and the JustRNA database covers 1,088,565 lncRNAs identified from 80 plant species^[13]^. However, the limitations of NGS, such as short read lengths and potential PCR amplification errors or bias pose great challenges for accurately identifying and quantifying lncRNAs^[15,16]^. In contrast, the third-generation-sequencing (TGS) is considered as a more suitable approach for studying lncRNAs due to its ability to generate long reads and single-molecules^[17]^. For instance, Wan et al. discovered about 28,000 lncRNAs in the mouse retina using the full-length isoform sequencing, with 3.4% of them belonging to intergenic lncRNAs (lincRNAs)^[18]^. Guan et al. identified 16,495 lncRNAs from 15,512 gene loci using Oxford Nanopore long-read sequencing in 19 chicken tissues^[19]^. They estimated that 70% of the new transcript originated from lncRNA loci, showing the value of long-read sequencing in discovering lncRNAs and resolving complex transcripts^[19]^. Recovering full-length lncRNA sequences facilitates a comprehensive understanding of their structure and functionality^[18,19]^.

Recent evidence has highlighted the ability of viruses to encode lncRNAs despite their small genomes^[20–24]^. Multiple viruses have been reported to encode lncRNAs, such as Kaposi’s Sarcoma-Associated Herpesvirus (KSHV), Human Cytomegalovirus (HCMV), and Human immunodeficiency virus (HIV). The virus lncRNAs (vlncRNAs) have been reported to play important roles throughout the viral life cycle^[21]^. For example, the vlncRNA PAN encoded by KSHV was found to activate viral replication and regulate viral gene expression^[21]^. The vlncRNA 2.7 encoded by the HCMV was reported to interact with complex I and prevents the translocation of GRIM-19, a gene associated with retinoid interferon-induced mortality, thereby maintaining the mitochondrial membrane potential and ultimately favoring viral ATP production^[22,23]^. Additionally, the vlncRNA ASP of HIV can recruit Polycomb Repressor Complex 2 to the 5’ LTR of HIV-1 to establish the latency of HIV-1^[24]^. These findings highlight the importance of vlncRNAs in regulating host cells during viral infections. Nevertheless, a comprehensive resource for vlncRNAs is still lacking, and the diversity, structure, and function of vlncRNAs remain largely unexplored. In this study, we systematically identified vlncRNAs for more than 20 viral species by mining viral infection-related TGS data and further characterized their structure and function. A database named vlncRNAbase was further built to store and organize these newly identified and known vlncRNAs. It would greatly facilitate further research on vlncRNAs in the field.

## Materials and methods

### The workflow of identifying vlncRNAs

The workflow for identifying vlncRNAs can be separated into three modules (Figure 1). The first module is Data collection, during which the viral infection-related TGS data generated by both Oxford Nanopore and PacBio SMRT technologies were obtained from NCBI GEO and SRA databases. The second module is Viral transcript identification, which includes identification of viral reads through three features: transcription start site (TSS), transcription end site (TES), and potential splice sites^[25]^, and filtering of viral transcripts by abundance and length. The third module is Viral transcript classification, which includes the identification of novel transcripts and the classification of viral transcripts into vmRNA or vlncRNA based on their coding potential.

**Figure 1.**
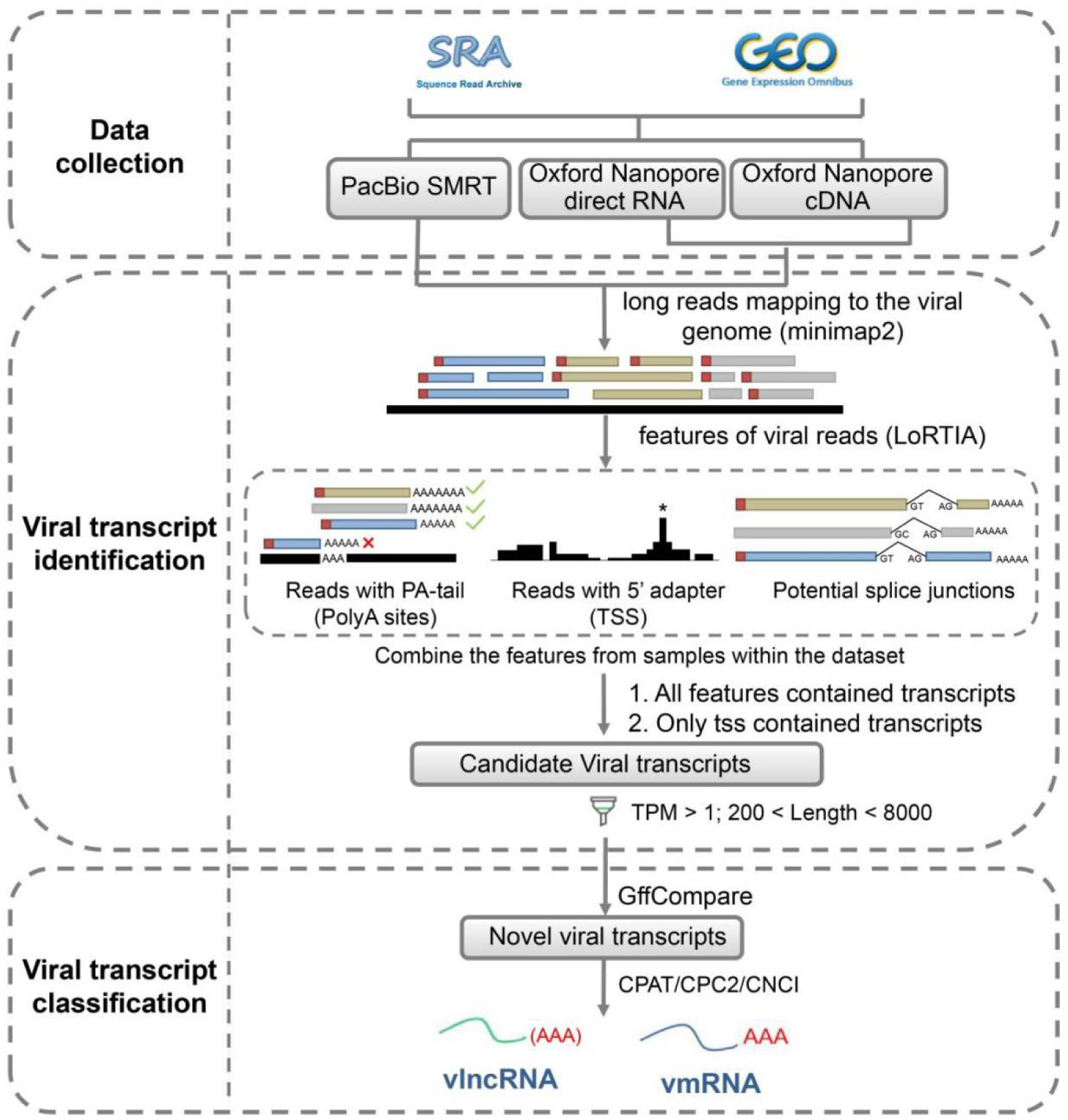
Workflow of identifying vlncRNAs. It contains three modules including Data collection, Viral transcript identification and Viral transcript classification.

### Data collection

The viral infection-related TGS datasets were collected from the NCBI SRA and GEO databases in June 2023. The keywords of “pacbio” AND “virus” or “nanopore” AND “virus” were used to query for viral infection-related TGS datasets from the two databases mentioned above. Then, only the datasets with the Library Source of “TRANSCRIPTOMIC” and Assay Type of “RNA-seq” were kept. Subsequently, the remaining datasets were inspected manually. A total of 36 TGS datasets, including 870 samples, were collected and used in the analysis, as provided in Table S1. The viral genomes used in these datasets and their related annotation files were manually collected from the NCBI GenBank database.

### Viral transcript identification

Firstly, viral transcripts were identified by mapping reads to viral genomes using minimap2 (version 2.17)^[26]^. The parameter settings differed depending on the sequencing type of the TGS data according to the official recommendation of minimap2: the parameter settings of “-ax splice:hq -uf”, “-ax splice -uf -k 14” and “-ax splice -k 14” were used for PacBio SMRT, Oxford Nanopore direct RNA, and Oxford Nanopore cDNA, respectively.

Second, to detect and annotate viral transcripts, we used LoRTIA (version 0.9.9)^[27]^ to analyze the mapped reads, using the following steps according to Fülöp’s study^[25]^: (1) The adapter and polyA sequences at the ends of the reads were checked to identify TSS and TES, respectively. (2) A Poisson distribution test was applied to the putative TSSs and TESs to eliminate random start and end sites, with Bonferroni correction for significance. (3) Potential splice junctions were identified, and those with common splice signals (GT/AG, GC/AG, AT/AC) were retained. Transcript features including TSS, TES and splice junctions were kept if they were detected in at least two reads and their read frequency exceeds 1‰ of all mapped reads to the viral genome. To improve accurate annotations, we combined features identified from different samples within the same dataset and filtered out viral transcripts supported by less than five reads. Finally, based on the accepted TSSs, TESs, and splice junctions, the sequences of these regions were combined to form viral transcripts. Since not all viral transcripts contain polyA sequences^[28]^, we also retained transcripts containing only TSS and determined their TES by selecting the longest read among mapped reads that had the same TSS. The transcripts with both similar TSS and TES (distance < 20 nt) were regarded as the same viral transcript, considering potential inaccuracies in the identification of TSS and TES. Then, the retained viral transcripts were then filtered by abundance and length. Only transcripts with abundance > 1 TPM (transcript per million) in at least one sample and length < 8000 nt were kept for further analysis.

### Viral transcript classification

Firstly, novel viral transcripts were identified using GffCompare (version 0.11.2)^[29]^. Only transcripts with class_code of “j”, “k”, “m”, “n”, which represent novel isoforms of known genes, and those of “i”, “x”, “u”, “y”, “o”, which potentially represent novel genes or transcriptional units^[30]^, were taken as novel transcripts and were provided in Table S2.

Then, three software tools including CPAT (version 3.0.4, parameter settings: “--antisense -- top-orf=5”)^[31]^, CPC2 (version 0.1, default parameter settings)^[32]^, CNCI (version 2, parameter settings: “-m ve”)^[33]^, were used to predict the coding potential of novel viral transcripts. If a transcript was predicted to be non-coding by at least two tools, or if it was predicted to be non-coding by a tool when there were only two predictions, it was considered as a lncRNA; otherwise, the mRNA.

### RNA family identification of vlncRNAs

To identify RNA families in vlncRNAs, firstly, the Rfam database (v14.9)^[34]^, a database of RNA families, was downloaded from https://ftp.ebi.ac.uk/pub/databases/Rfam/14.9/. Then, the cmscan program from Infernal (v1.1.2)^[35]^ with parameter settings of “--rfam --nohmmonly -- fmt 2 --cut_ga” was used to identify RNA families in the Rfam database based on vlncRNA sequences. All RNA family annotation results for vlncRNAs were summarized in Table S3.

### Analysis of host RNA mimicry of vlncRNAs

Since most TGS data were related to human viruses, we focused on the human RNA mimicry of vlncRNAs. Firstly, sequences of human lncRNAs were collected from Ensemble (release-109, https://ftp.ensembl.org/pub/release-109/fasta/homo_sapiens/ncrna/). Then, the MMseqs2 (version 6.f5a1c)^[36]^ with the easy-cluster mode and with parameter setting of “--min-seq-id 1 --cov-mode 0” was used to remove sequence redundancy at 100% level for both human lncRNAs and novel vlncRNAs. The sequence similarity between human lncRNAs and novel vlncRNAs was analyzed using BLASTN (version 2.9.0+, with the “-word_size 11” parameter). The vlncRNA was considered to be similar to human lncRNAs if the e-value was smaller than 1E-5 in the blast results. The secondary structure of human lncRNAs was predicted using LinearFold (version 1.0). Subsequently, the RNAdistance (version 2.6.4)^[37]^ with default parameters from the ViennaRNA package was employed to calculate the distance, i.e., the fscore (the lower fscore indicates higher structure similarity), between a pair of human lncRNAs and vlncRNAs. The fscores between all pairs of human lncRNAs and vlncRNAs were ranked in an increasing order. The 5% percentage of these fscores (329) was taken as the threshold, which is much stricter than those used by previous studies ^[38–40]^ such as the lower quartile of fscores used by Leung^[39]^. A pair of human lncRNA and vlncRNAs was considered similar if the distance between them was smaller than the threshold. RNA secondary structures were visualized by forna in the ViennaRNA package^[37]^.

### Predicting interactions between human miRNAs and virus-like hlncRNAs as well as human-mimicry vlncRNAs

A total of 1,917 human miRNAs were obtained from miRBase (v22, https://www.mirbase.org/)^[41]^. The interactions between human miRNAs and virus-like hlncRNAs as well as human-mimicry vlncRNAs were predicted using miRanda (version 3.3a)^[42]^. To minimize the false-positive rate of the method^[43]^, a strict criterion was used: only a seed region complementarity score (sc) greater than 200, and energy less than −40 kcal/mol were used for further analysis.

### Functional analysis of human lncRNAs and human miRNAs

The Gene Ontology (GO) and Kyoto Encyclopedia of Genes and Genomes (KEGG) enrichment analysis was conducted to reveal the functions of hlncRNAs and miRNAs using the *enrichGO()* and *enrichKEGG()* functions in the clusterProfiler package (version 4.8.3)^[44]^. GO terms and KEGG pathways with FDR adjusted p-value less than 0.05 were considered significantly enriched.

### Analysis of the alternative splicing of vlncRNAs

To investigate the alternative splicing of vlncRNAs and vmRNAs, SUPPA (version 2.3)^[45]^ with parameters settings of “generateEvents -f ioe -e SE SS MX RI FL” was used to identify the type of splicing events.

### Experimental validation of vlncRNAs

Cells: the Madin-Darby canine kidney (MDCK) cell line, HEK293T cells, and Vero cells were purchased from ATCC. The Huh7 cell line was kindly shared by Haizhen Zhu (Hunan University, Changsha, China). All cells were propagated in Dulbecco’s modified Eagle medium (DMEM) supplemented with 10% Fetal Bovine Serum (FBS), and 1% penicillin-streptomycin. Cells were maintained in a humidified atmosphere of 5% CO_2_ at 37°C.

Viruses: the virus Indiana vesiculovirus (VSV) was kindly shared by Haizhen Zhu (Hunan University, Changsha, China). The strain A/California/07/2009 of Influenza A virus (IAV) and VSV were propagated and amplified at MOI of 0.001 in chicken embryos and HEK293T cells, respectively. Supernatants were harvested and clarified by centrifugation, then stored at −80 °C. Viral titers were determined by plaque assay in Vero cells.

Real-time PCR assay: then, utilizing the aforementioned amplified viruses with MOI=0.5, Huh7 cells were infected with VSV and cultured for an additional 9 hours, while MDCK cells were infected with IAV and cultured for an additional 24 hours. Subsequently, the total cellular RNA was isolated using TRIzol Reagent (Invitrogen). After the RQ1 DNase (Promega) treatment, the extracted RNA was used for the synthesis of the first strand of cDNA with the Superscript III first-strand Synthesis System (Invitrogen) as described previously^[28,46]^. After the reverse transcription (RT), the generated template cDNAs were utilized for real-time quantitative PCR (qPCR) analysis. To reduce non-specific amplification, qPCR primers were designed to strategically target alternative splice sites in vlncRNAs. The primers used for RT-qPCR are listed in Table S4. Expression levels of vlncRNAs were normalized by mRNA of the housekeeping gene GAPDH. The relative expression levels of vlncRNAs were calculated using the method of 2-^△△^ct.

### Database implementation

The front-end and back-end separation frameworks were used to implement vlncRNAbase which was described in our previous study^[47]^. The JavaScript Object Notation format was employed to implement the communication between the client-side and server-side layers. The data were stored in a MySQL (version 5.5.62) database^[47]^.

### Statistical analysis

All statistical analyses were conducted in Python (version 3.8) and R (version 4.3.1). The Wilcoxon rank-sum test was conducted using the function of *wilcox.test()* in R. A p-value of less than 0.05 was considered statistically significant.

## Results

### Identification and validation of novel vlncRNAs from TGS data

A total of 5,053 novel vlncRNAs and 7,101 novel vmRNAs were identified in 25 virus species from 17 virus families that belonged to 4 Baltimore groups based on computational analysis of 870 viral infection-related samples of TGS data (Materials and Methods) (Figure 2A & Figure S1). Most novel vlncRNAs were identified from dsDNA viruses (97.94%), then the ssRNA (+) (1.39%) and ssRNA (−) (0.79%). The Vaccinia virus (VACV) of the *Poxviridae* family had the highest number of novel vlncRNAs (1766), followed by Human betaherpesvirus 5 (HHV-5) (749) and Monkeypox virus (MPXV) (627). Similarly, most novel vmRNAs were also identified from double-stranded DNA (dsDNA) viruses (95.40%). The HHV-5 of the *Herpersviridae* family, Haliotid herpesvirus 1 (HaHV-1), and Ostreid herpesvirus 1 (OsHV-1) from the *Malacoherpesviridae* family had the largest number of novel vmRNAs, with 1433, 1171, and 1088, respectively. The number of novel vmRNAs exceeded that of novel vlncRNAs for all viruses except VACV, MPXV, Human alphaherpesvirus 3 (HHV-3), African swine fever virus (ASFV), and West Nile virus (WNV).

**Figure 2.**
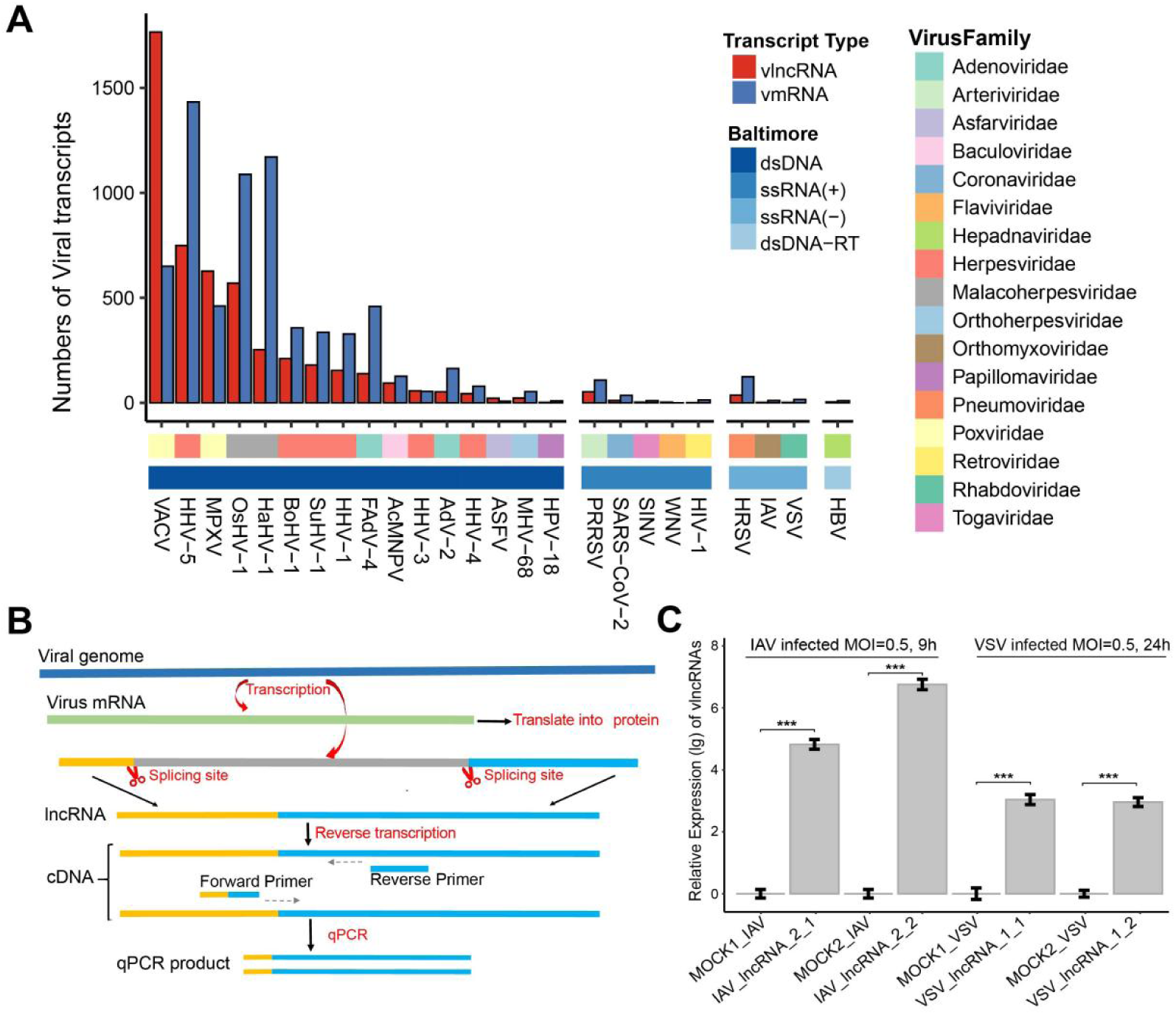
Identification and characterization of novel vlncRNAs and vmRNAs. (A) The number of novel vlncRNAs (red) and vmRNAs (blue) in 25 virus species. The first bar below the x-axis represents the Baltimore group to which the virus belongs, while the second bar represents the virus family. For clarity, the abbreviation of virus names was used. Supplementary Table S5 displays the full names of these viruses. dsDNA, double-stranded DNA virus; ssRNA (+), positive-sense single-stranded RNA virus; ssRNA (−), negative-sense single-stranded RNA virus; dsDNA-RT, double-stranded DNA reverse transcribing virus. (B) Diagram of designing primers specific for detecting vlncRNAs by RT-qPCR. (C) The relative expression of IAV_lncRNA_2 and VSV_lncRNA_1 by RT-qPCR following amplification with two pairs of primers. For clarity, the expression values were log10 transformed. ***: p-value < 0.001.

To validate the vlncRNAs, two vlncRNAs of IAV, IAV_lncRNA_1 from the M1 segment and IAV_lncRNA_2 from the PB1 segment, and two vlncRNAs of VSV, VSV_lncRNA_1 from the gp2 gene and VSV_lncRNA_2 from the gp1 gene, were selected for validation by RT-qPCR (Materials and Methods). To ensure that the primers specifically amplify the target fragments, we designed two pairs of primers targeting the alternative splice sites of each vlncRNA (Figure 2B & Table S4). Subsequently, RT-qPCR was used to quantify the relative expression levels of these vlncRNAs. Two vlncRNAs, i.e., the IAV_lncRNA_2 and VSV_lncRNA_1, exhibited high expression levels at 9 hours post-infections (hpi) and 24 hpi, respectively. The former showed relative fold changes of 4.82 and 6.76 for two pairs of primers after log_10_ transformation, while the latter had 3.04 and 2.96 for two pairs of primers after log_10_ transformation (Figure 2C). In contrast, almost no expression was detected in the mock-infected group.

### Length, GC content and abundance of vlncRNAs

Then, the length, GC content and abundance of vlncRNAs and vmRNAs were analyzed and compared (Figure 3A-C). The vlncRNAs were found to have significantly shorter length, lower GC content, and lower expression level compared to those of vmRNAs. Specifically, in terms of length, vmRNAs were twice as long as vlncRNAs, with the former having a median of 2,020 nt and the latter having a median of 876 nt. Concerning the GC content, vmRNAs had a median of 48.27%, while vlncRNAs had a median of 37.40%. Regarding the abundance, vmRNAs and vlncRNAs had median log_2_TPM values of 6.35 and 5.54, respectively.

**Figure 3.**
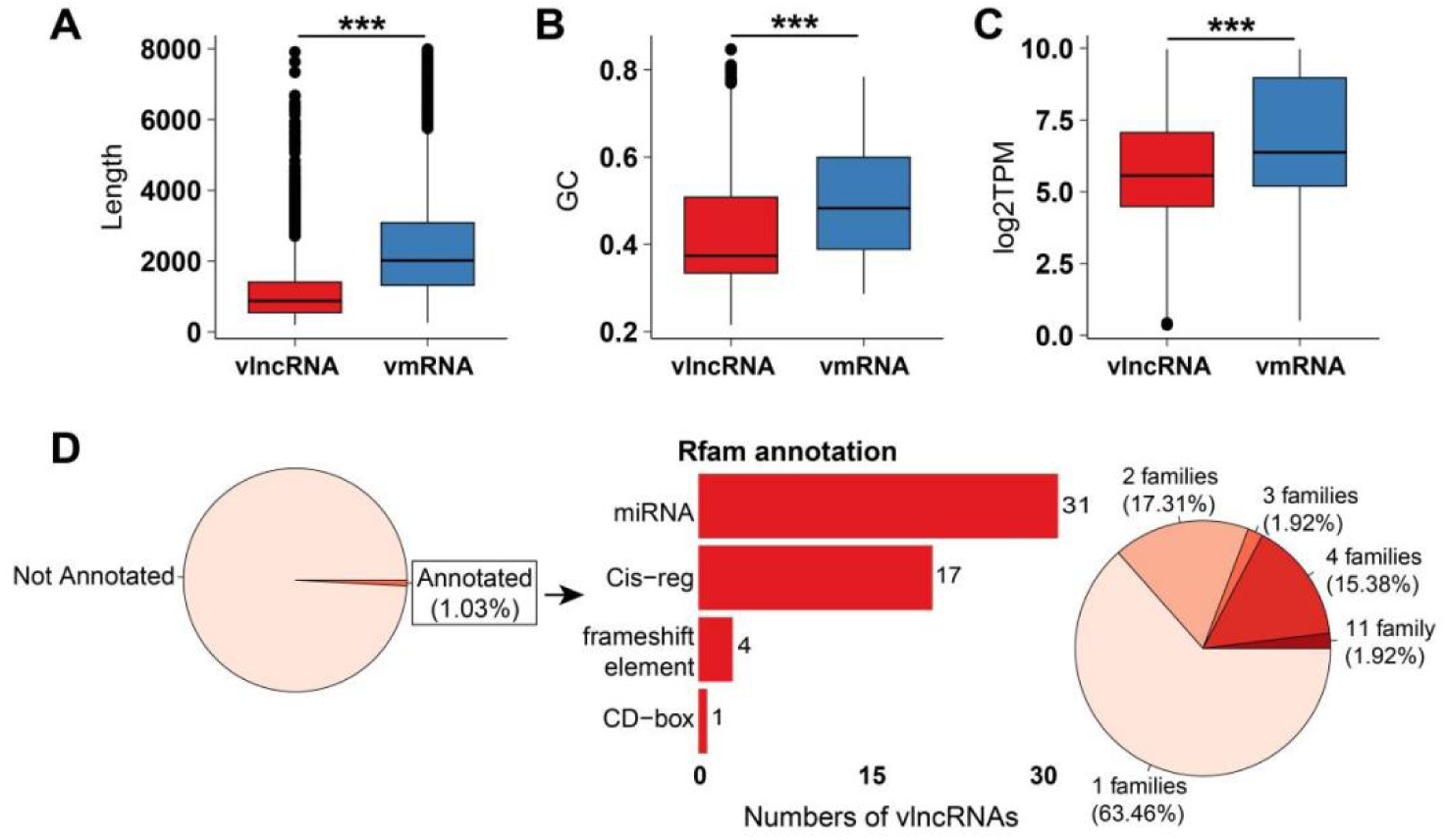
Sequence and structure characteristics of novel vlncRNAs. (A)∼(C) refer to the length, GC content and expression distribution of novel vmRNAs and vlncRNAs, respectively. For clarity, the expression of RNAs were log2-transformed. (D) The pie chart at left refers to the portion of the vlncRNAs annotated in Rfam. The bar plot in the middle shows the number of vlncRNAs that were classified into four RNA types based on the Rfam. The pie chart at right shows the proportion of vlncRNAs that were annotated to one or multiple RNA families.

### Analysis of the structural characteristics of novel vlncRNAs

Then, RNA families of the vlncRNAs were annotated in the to further characterize their structure based on the Rfam database (Materials and Methods). Only 52 vlncRNAs (1.03% of all vlncRNAs) were annotated and predicted to contain 1∼11 RNA families. Among them, 63.46% had only one annotated RNA family (Figure 3D). A total of 33 RNA families were identified in vlncRNAs and they were further grouped into four RNA types including “miRNA” that is defined as small noncoding and single-stranded RNAs involved in the gene expression regulation, “Cis-reg” that is defined as functional regulatory elements to regulate the transcription of neighboring genes, “frameshift element” that is defined as a sequences that facilitate ribosomal frameshifting during translation, and “CD-box” that was defined as snoRNA with C (UGAUGA) and D (CUGA) box motifs. “miRNA” was the most frequently annotated RNA type, with 31 vlncRNAs; “Cis-reg” was the secondly annotated RNA type, containing 17 vlncRNAs. The other two RNA types each contained less than five vlncRNAs.

### Extensive structural mimicry of human lncRNAs by vlncRNAs

Previous studies have shown that viral RNAs such as miRNAs, may mimic the host RNAs to suppress innate immunity and hijack host machinery for their survival^[48]^. Thus, the sequence and structure similarity between vlncRNAs and human lncRNAs (hlncRNAs) was analyzed, as most viruses analyzed were human-infecting. Only 0.14% of vlncRNAs had sequence similarity with hlncRNAs (Materials and Methods), suggesting little sequence similarity between vlncRNAs and hlncRNAs. Then, the similarity of secondary structures between all pairs of vlncRNAs and hlncRNAs was analyzed (Materials and Methods) (Figure 4A). A median fscore (smaller fscore indicates higher similarity) of 906 was observed between vlncRNAs and hlncRNAs. For comparison, we randomly selected 1,000 hlncRNAs and calculated the structure similarity among them. A median fscore of 1,023 was obtained for hlncRNAs, which was significantly larger than that between vlncRNAs and hlncRNAs (p-value < 2.2E-16 in the wilcox rank-sum test). This suggests that vlncRNAs had greater structural similarity to hlncRNAs than the hlncRNA do with each other. The top 5% pairs between vlncRNAs and hlncRNAs were considered as structural similar, which involved 772 vlncRNAs (defined as human-mimicry vlncRNAs) from 15 human viruses and 9,393 hlncRNAs (defined as virus-like hlncRNAs).

**Figure 4.**
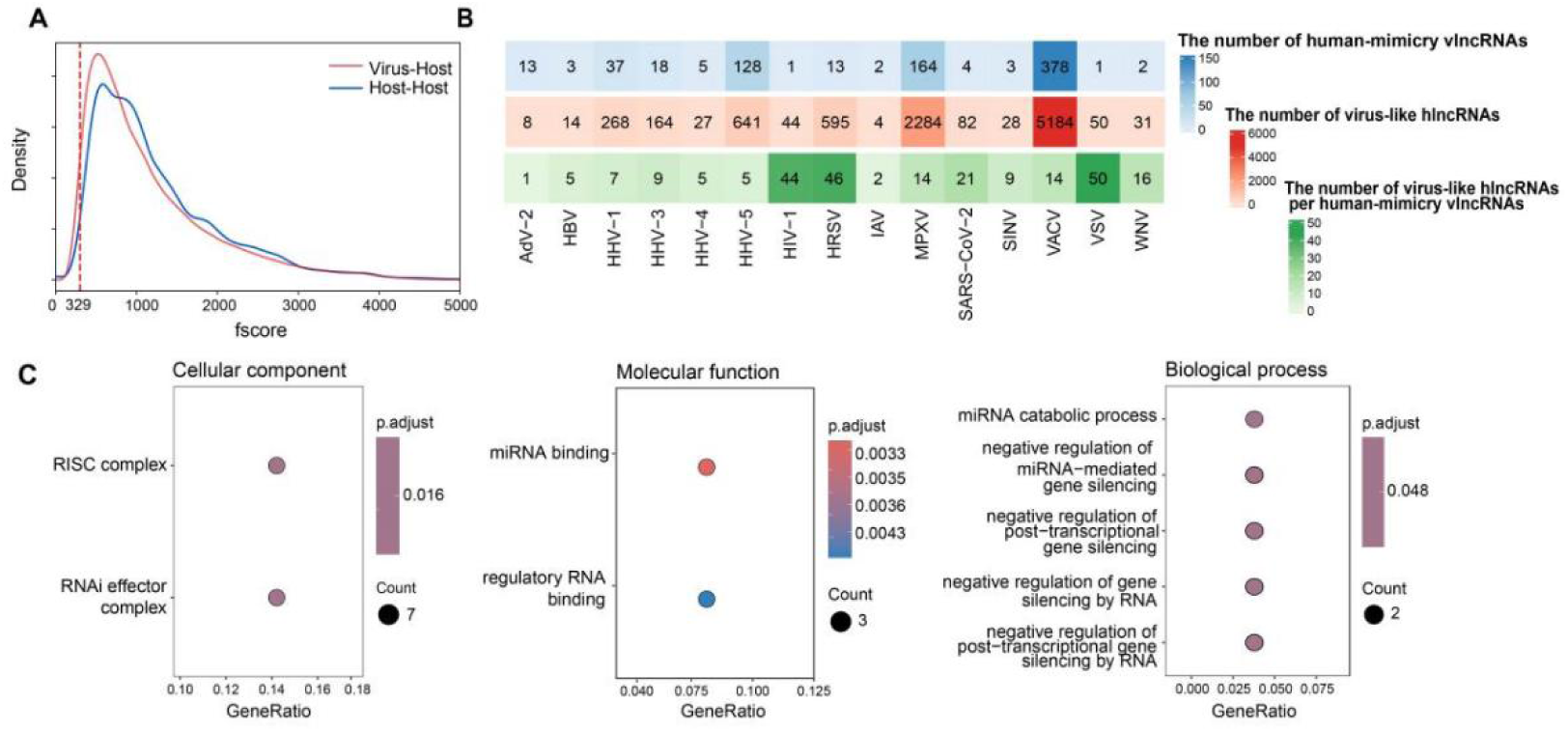
Analysis of hlncRNAs mimicry of vlncRNAs. (A) The density distribution of fscores between vlncRNAs and hlncRNAs, as well as among hlncRNAs. The dashed line represents the 5% threshold for the similarity between vlncRNAs and hlncRNAs. (B) The number of human-mimicry vlncRNAs and virus-like hlncRNAs across 15 human viruses. The first row (blue) displays the number of human-mimicry vlncRNAs, the second row (red) presents the number of virus-like hlncRNAs, and the third row (green) displays the number of virus-like hlncRNAs per human-mimicry vlncRNAs. (C) Enriched GO functions (Cellular Component, Molecular Function and Biological Process) for virus-like hlncRNAs that were structurally similar to vlncRNAs of at least 8 viruses.

Human viruses exhibited a range of 1-378 human-mimicry vlncRNAs, with a median of 5. VACV had the highest number of human-mimicry vlncRNAs (378). On the other hand, the number of virus-like hlncRNAs also varied across viruses, ranging from 8 to 5184. VACV had the largest number of virus-like hlncRNAs (5,184), followed by MPXV (2,284) and HHV-5 (641). We further calculated the number of virus-like hlncRNAs per human-mimicry vlncRNAs (Figure 4B). Interestingly, the VSV had the highest number of virus-like hlncRNAs per human-mimicry vlncRNAs (50), followed by Human Respiratory Syncytial Virus (HRSV) (46) and Human alphaherpesvirus 1 (HHV-1) (44).

Then, the functions of vlncRNAs were investigated by analyzing the functions of virus-like hlncRNAs, as similar structures may suggest similar functions. A total of 1466 virus-like hlncRNAs were found to be structurally similar to vlncRNAs in at least half of viruses analyzed (8 out of 15). These virus-like hlncRNAs were enriched in a limited number of Gene Ontology (GO) terms (Figure 4C) and showed no significant enrichment in the KEGG pathway. Specifically, the enriched cellular components included “RISC complexes” and “RNAi effector complexes”; the enriched molecular functions included “miRNA binding” and “regulatory RNA binding”; the enriched biological processes were primarily associated with the negative regulation of gene silencing, such as “negative regulation of miRNA-mediated gene silencing” and “negative regulation of post-transcriptional gene silencing”.

### Human-mimicry vlncRNAs and virus-like hlncRNAs binding to similar miRNAs

The lncRNAs can bind to miRNAs to take part in post-transcriptional processes^[7]^. Our results showed that the virus-like hlncRNAs were mainly involved in miRNA-related functions (Figure 4C). Thus, we further investigated whether the human-mimicry vlncRNAs and their corresponding virus-like hlncRNAs bind to similar miRNAs. The analysis showed that nearly all human-mimicry vlncRNAs (768/772) bound to similar miRNAs as their corresponding virus-like hlncRNAs. The number of shared miRNAs between human-mimicry vlncRNAs and virus-like hlncRNAs ranged from 0 to 184, with a median of 37 and a mean of 42. For instance, the VACV_lncRNA_117 and its corresponding virus-like hlncRNA ENST00000457058.1 had highly similar structures (Figure 5B). They interacted with 48 and 51 human miRNAs, respectively. Of these, six miRNAs bound to both lncRNAs (colored in red in Figure 5B).

**Figure 5.**
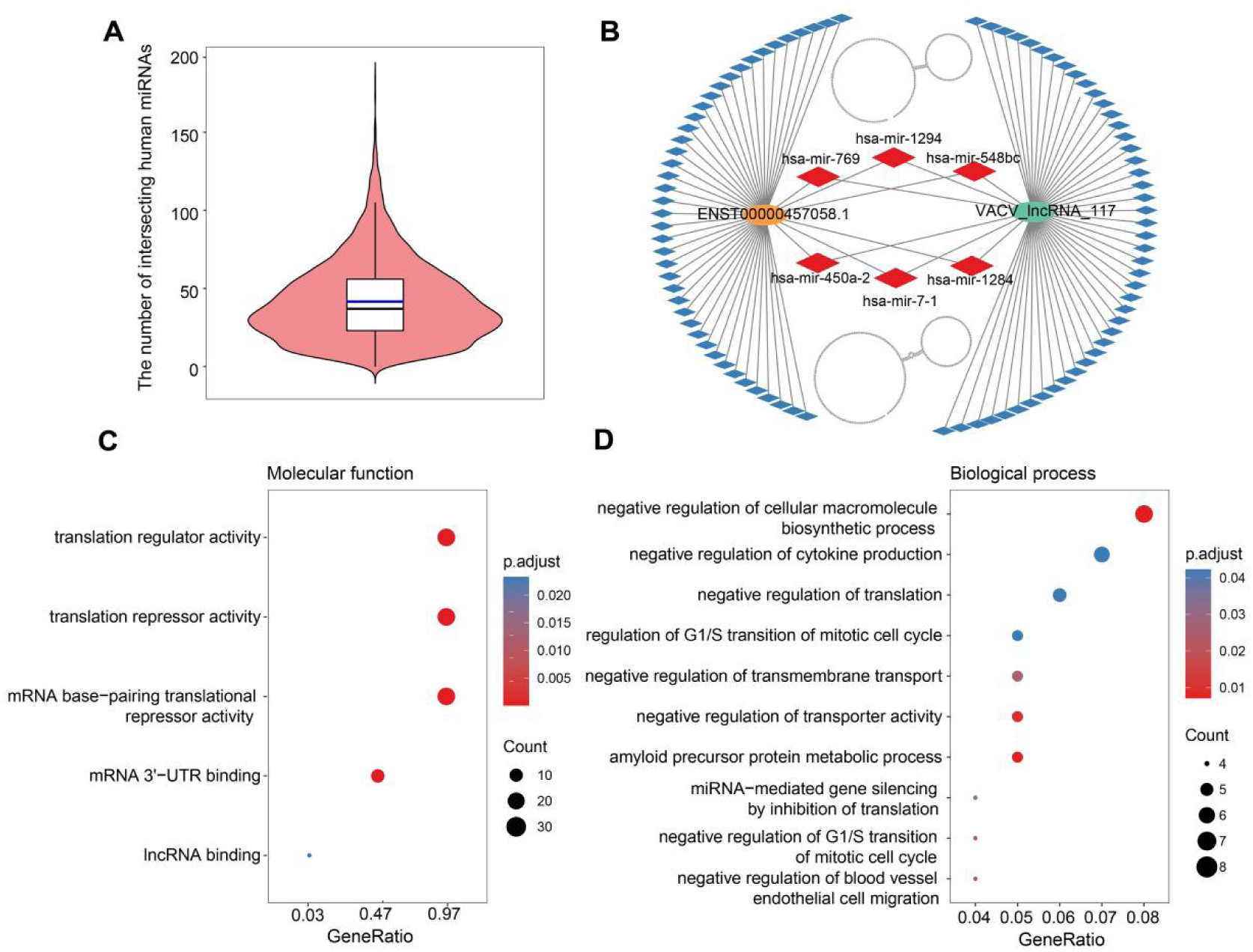
Analysis of miRNAs interacting with both human-mimicry vlncRNAs and virus-like hlncRNAs. (A) The distribution of the number of human miRNAs that interacted with both the human-mimicry vlncRNAs and their corresponding virus-like hlncRNAs. (B) An example of a pair of human-mimicry vlncRNAs and virus-like hlncRNAs binding to similar miRNAs. The secondary structures of the hlncRNA (top) and vlncRNA (bottom) are shown. miRNAs are represented by diamonds, with red ones indicating miRNA that interact with both hlncRNA and vlncRNA. (C)∼(D) Enriched molecular functions and biological process for human miRNAs that interact simultaneously with virus-like hlncRNAs and human-mimicry vlncRNAs.

The functions of 505 human miRNAs that interacted simultaneously with virus-like hlncRNAs and human-mimicry vlncRNAs were investigated by the GO and KEGG pathway enrichment analysis. A limited number of functions were enriched. The enriched molecular functions were mainly related to translation regulation, such as “translation regulator activity” and “mRNA base-pairing translational repressor activity” (Figure 5C). The enriched biological processes were mainly related to negative regulation of multiple essential life processes such as cellular macromolecule biosynthesis, cytokine production, translation, transmembrane transport and regulation of cell cycle such as G1/S transition of the mitotic cell cycle (Figure 5D). The enriched cellular components included the RISC complex and RNAi effector complex pathways, similar to those for virus-like hlncRNAs (Figure S2A). The enriched KEEE pathway included the “microRNA in cancer” pathway (Figure S2B).

### Expression correlation between human-mimicry vlncRNAs and virus-like hlncRNAs

Then, the relationship between the expression of human-mimicry vlncRNAs and corresponding virus-like hlncRNAs was investigated. The analysis revealed a Pearson correlation coefficient of 0.20 (p-value < 0.001) (Figure S3), indicating a weak but statistically significant positive correlation between the expression level of these two RNA types.

### Generation dynamics of vlncRNAs

To explore the expression dynamics of vlncRNA during viral infections, we analyzed the number of vlncRNAs generated at different time points post-infection across eight viruses. For most of these viruses, the number of vlncRNAs per sample increased as viral infections progressed (Figure 6A and Figure S4). For example, in MPXV, the number of vlncRNAs per sample increased from 49 at 1 hpi to 74 at 4 hpi, and further to 98 at 12 hpi. Similarly, in VACV, the number of vlncRNAs per sample increased from 23 at 1 hpi to 26 at 3 hpi, and further to 32 at 8 hpi (Figure 6A). Some viruses, including HHV-1, HHV-4, and SuHV-1 showed complex vlncRNA expression patterns. For example, HHV-4 maintained a relatively stable capacity to generate vlncRNAs throughout the infection period (from 10 min to 72 hpi), although some fluctuations.

**Figure 6.**
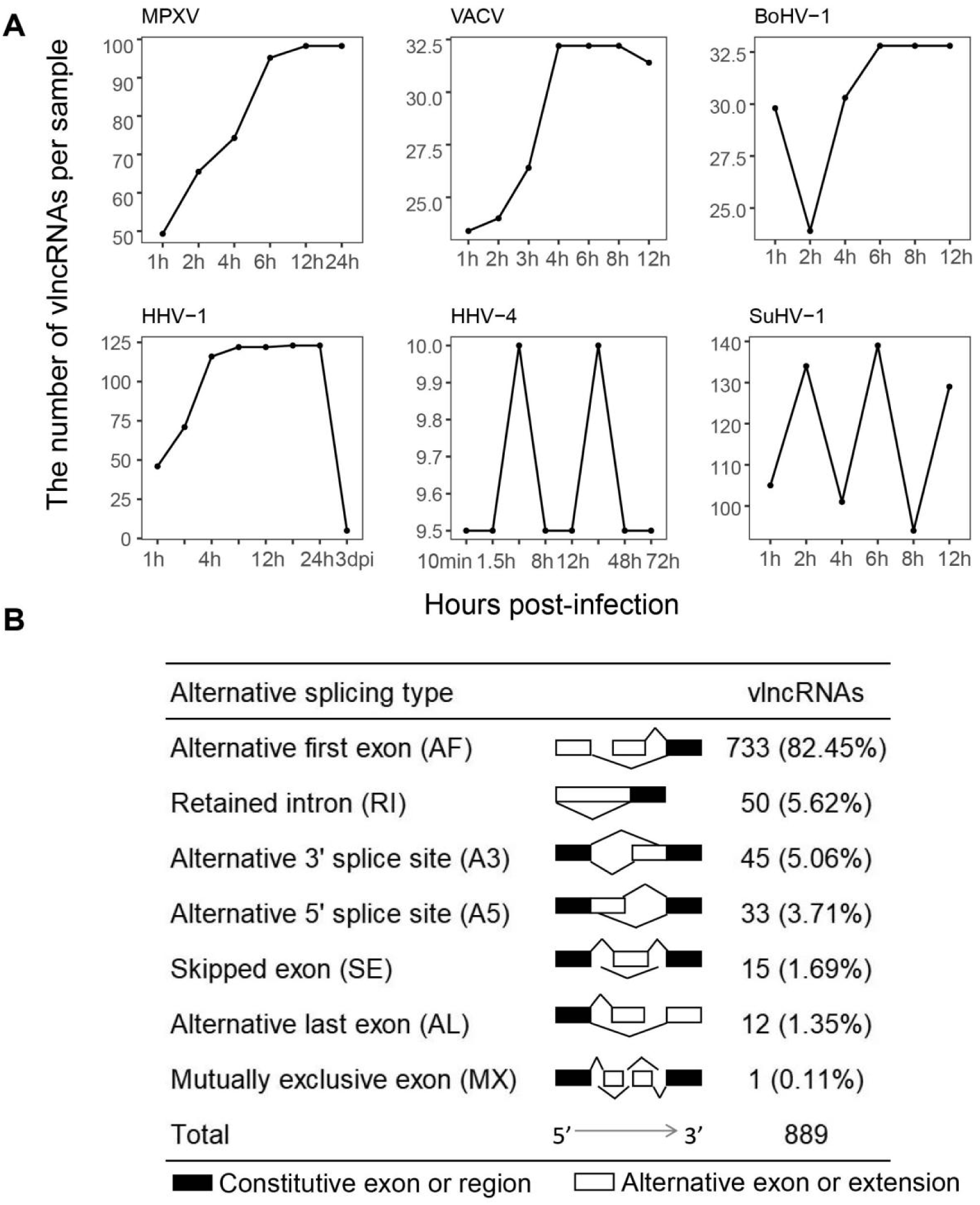
Analysis of vlncRNA generation dynamics and the biogenesis mechanism. (A) The number of vlncRNAs per sample detected at different hours post-infection in Monkeypox virus (MPXV), Vaccinia virus (VACV), Bovine alphaherpesvirus 1 (BoHV-1), Human alphaherpesvirus 1 (HHV-1), Human gammaherpesvirus 4 (HHV-4), Suid alphaherpesvirus 1 (SuHV-1). (B) Analysis of the alternative splicing pattern of vlncRNAs.

### Analysis of the alternative splicing patterm of vlncRNAs

To investigate the biogenesis mechanism of vlncRNAs, we analyzed seven common alternative splicing (AS) patterns of vlncRNAs. The results showed that the alternative first exon (AF) events accounted for more than 80% of the total AS events (Figure 6B), a splicing pattern similar to that observed in humans and mice^[49]^, where AF events also dominate; Retained intron (RI) and Alternative 3’ splice site (A3) both accounted for more than 5% of the total AS events, while the remaining types including Alternative 5’ splice site (A5), Exon skipping (SE), Alternative last exon (AL), and Mutually exclusive exons (MX) each accounted for less than 5% of AS events. Interestingly, the AS patterns of vlncRNAs differed between enveloped and non-enveloped viruses. In enveloped viruses, AF accounts for 91.07%, followed by RI at 3.92%, while in non-enveloped viruses, three AS events with proportions exceeding 10% including AF (40.00%), RI (24.67%), and A3 (17.33%) (Table S6).

### Construction of a vlncRNA database

Finally, a database named vlncRNAbase was created to store and organize the vlncRNAs identified above. Besides, 15 vlncRNAs experimentally-validated by PCR-based methods (RT-qPCR, RACE-PCR, etc.) and Sanger sequencing, and 1,089 vlncRNAs that have been identified through high-throughput methods were curated from public databases including NCBI RefSeq, GenBank and RNAcentral, and were also provided in the database. The vlncRNAbase is freely available to the public at http://computationalbiology.cn/vlncRNAbase/#/. It mainly includes pages of Browse, Search, Expression, Interaction, Statistic, Download, and Help which were described briefly as follows (Figure 7).

**Figure 7.**
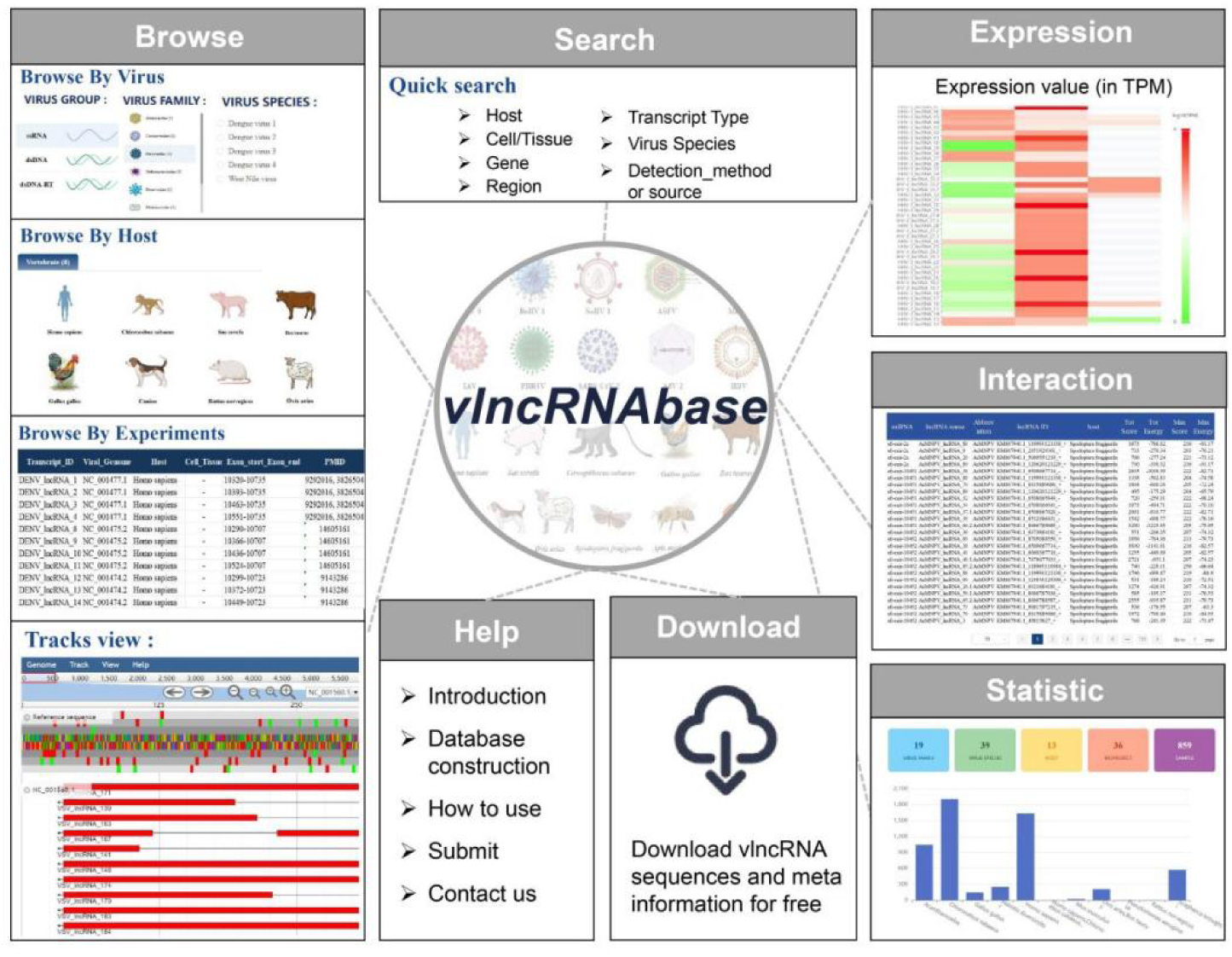
The structure of the vlncRNAbase. It mainly includes pages of Browse, Search, Expression, Interaction, Statistic, Download, and Help.

#### Browse

This page displays vlncRNAs by virus, host or experiments. When browsing by virus, a virus of interest should be selected, and the vlncRNAs identified for the virus were provided in a table. The details about the vlncRNAs such as strand, abundance, sequence, and detection method or source are provided. These vlncRNAs can also be visualized in a genome browser. When browsing the vlncRNAs by host, a host of interest is first selected; then, viruses infecting the host are provided in a table. The vlncRNAs identified for these viruses can be visualized as above. When browsing the vlncRNAs by experiment, all experimentally-validated vlncRNAs are provided in a table.

#### Expression

This page displays the expression values of vlncRNAs in TGS samples. Firstly, a bioproject is selected; then, the expression values of vlncRNAs identified in the bioproject are shown in a heatmap. The Run accession number, the cell or tissue of the sample, the expression value, and the vlncRNAs ID appear in a small window when hovering over the heatmap.

#### Search

Users can search for vlncRNAs of interest by virus, host, cell or tissue, gene, genomic regions, and detection method or source.

#### Interaction

The page displays the interactions between host miRNAs and vlncRNAs predicted by miranda (version 3.3a).

#### Statistic

This page displays a summary statistic about the number of vlncRNAs by virus host and the length distribution of vlncRNAs for each virus.

#### Download

The details of vlncRNAs stored in the database are freely available for downloading by virus which is organized by virus family.

#### Help

This page displays the method about the data curation process and the tutorial of using vlncRNAbase.

## Discussion

The vlncRNA has been reported to play important regulatory roles in virus infections^[20,22–24]^. Although an increasing number of vlncRNAs have been reported in recent years, most of these studies have focused on one or a few vlncRNAs, or vlncRNAs from one or a few viruses. This study for the first time systematically identified and characterized vlncRNAs from over 20 viral species by mining the viral infection-related TGS data, which provides a valuable resource for further study of vlncRNAs. Additionally, the structure, function, dynamics and biogenesis mechanism of vlncRNAs were further investigated, which deepens our understanding of vlncRNAs.

Previous studies have shown that the ability of different viruses to encode RNA products varies widely, including circRNAs^[50,51]^ and sRNAs^[47,52]^. This study found that the ability of encoding vlncRNAs also varied much among viruses, with most vlncRNAs identified from dsRNA viruses, followed by ssRNA (+) viruses. Interestingly, viruses of the *Herpesviridae* family were found to encode a large number of vlncRNAs (Figure 2A), likely due to their large genome size. Although two novel vlncRNAs were experimentally validated in the study, further efforts are needed to verify the existence of vlncRNAs and clarify their functions.

Viral RNAs have been reported to mimic host RNAs to exert their functions^[48]^. For example, miR-K12-11 of the KSHV can mimic host miR-155 to attenuate transforming growth factor beta (TGF-β) signaling, thus facilitating viral infection and tumorigenesis^[48]^. Unfortunately, there have been no reports about the mimicry of host lncRNAs by vlncRNAs. This may be due to the difficulty in identifying sequence similarity between lncRNAs since the lncRNAs lack conserved seed regions as miRNAs. Surprisingly, a large number of vlncRNAs were observed to be structurally similar to hlncRNAs. Among them, vlncRNAs encoded by VACV showed the strongest mimicry of hlncRNAs. Considering that the VACV stably generated vlncRNAs throughout the infection period, vlncRNAs may play an important role in viral infection.

The lncRNA plays an important role in translation regulation by binding to miRNAs^[8]^. Our results showed that virus-like hlncRNAs were mainly involved in the negative regulation of gene silencing, possibly by binding to miRNAs. Interestingly, the human-mimicry vlncRNAs shared a significant number of miRNAs with the corresponding virus-like hlncRNAs and these shared miRNAs were mainly involved in negative regulation of essential life processes. This suggests that the vlncRNA may action as a miRNA sponge, similar to the role of circular RNA, by mimicking the structure of host lncRNAs. Both the vlncRNA and human lncRNA cooperatively inhibit the function of miRNA and promote the essential life process of the host cell, which may facilitate the proliferation of the virus (Figure 8).

**Figure 8.**
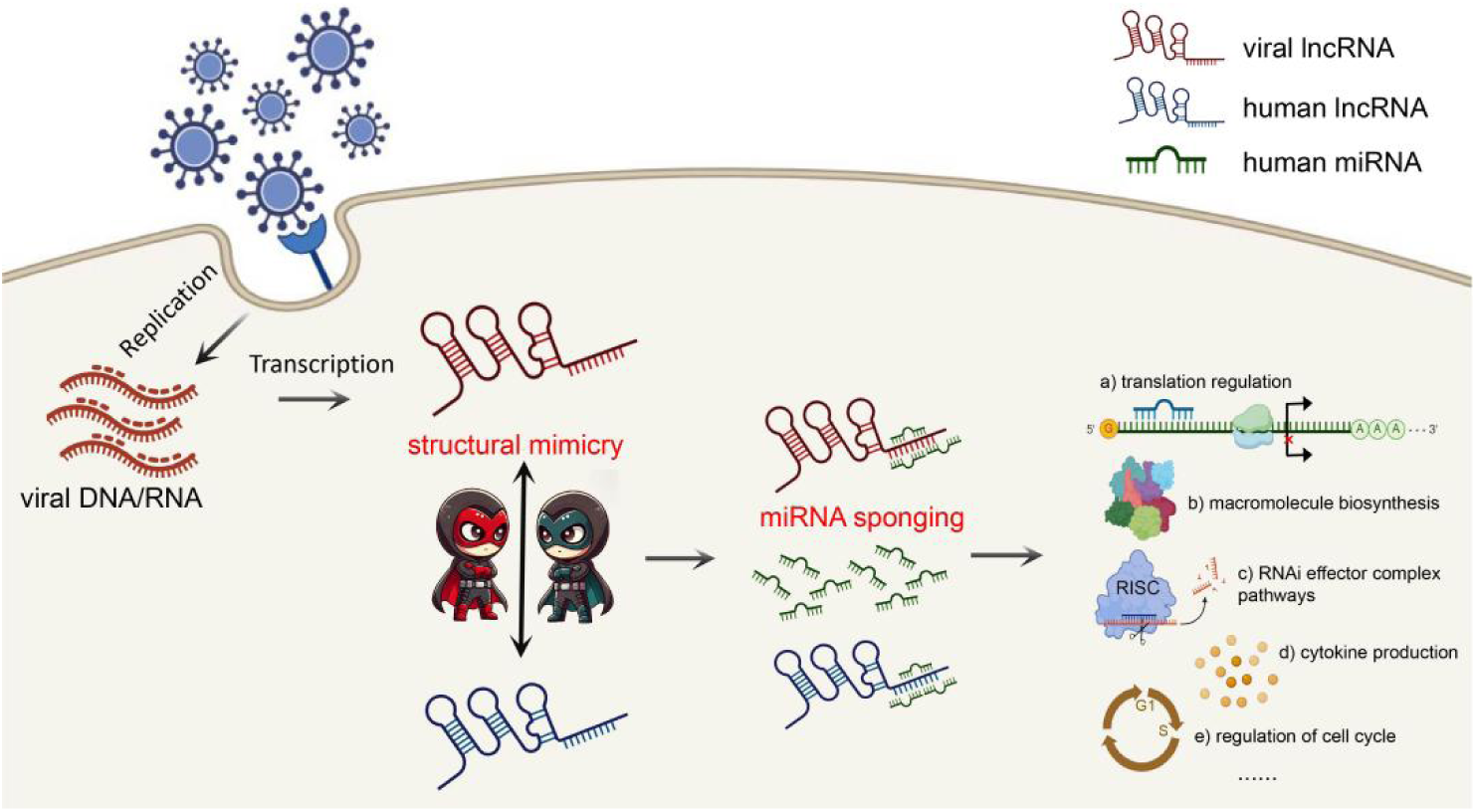
The hypothesized mechanism of how vlncRNAs function in the host cell. The vlncRNA may act as miRNA sponge by mimicking the hlncRNA structures, which cooperates with hlncRNAs to inhibit miRNA function and promote the essential life process of the host cell, thus facilitating viral proliferation.

This study has several limitations. Firstly, for positive-strand RNA viruses, it is difficult to distinguish between subgenomic RNAs and lncRNAs, although the subgenome has some features such as the leader transcription regulatory sequence^[53]^. Secondly, despite the inclusion of over 20 viruses, the list is still small. Besides, the samples used in the study have a bias towards some viruses such as herpesviruses, requiring periodic updates to capture the diverse landscape of novel vlncRNAs in more viruses. Lastly, the functions of vlncRNAs may have been overlooked for a long time. Much more effort is needed to clarify the function and the biogenesis mechanism of vlncRNAs.

## Conclusion

Overall, this study systematically identified and characterized a large number of vlncRNAs from more than 20 virus species for the first time. We found that the vlncRNAs may act as a miRNA sponge by structurally mimicking the host lncRNAs. The study deepens our understanding of the diversity and complexity of vlncRNAs and provides a valuable resource for further studies. Besides, it provides new insights into the structure and function of the vlncRNAs.

## Availability of data and materials

All data used in the study are available in supplementary materials. The vlncRNAbase is publicly available at http://computationalbiology.cn/vlncRNAbase/#/.

## Authors’ contributions

Ping Fu and Zena Cai analyzed the data. Ping Fu wrote the draft and built the vlncRNAbase; Ruina You performed the RT-qPCR experiment; Lei Deng and Zhaoyong Li supervised the experiment and revised the manuscript; Zhichao Miao revised the manuscript and provided part of funding for the project; Yousong Peng designed and supervised the project, wrote and revised the manuscript.

## Funding

This work was supported by the R&D Program of Guangzhou National Laboratory (GZNL2024A01002), National Natural Science Foundation of China (32170651 & 32370700), and Hunan Provincial Natural Science Foundation of China (2024JJ2015).

## Acknowledgements

We thank Prof. Haizhen Zhu in Hunan University for kindly providing the VSV strain and the Huh7 cell line. We thank members in PengLab for helpful discussions on the manuscript.

## Competing interests

The authors declare that they have no competing interests.

## Notes

### Competing Interest Statement

The authors have declared no competing interest.

